# Mining Thousands of Genomes to Classify Somatic and Pathogenic Structural Variants

**DOI:** 10.1101/2021.04.21.440844

**Authors:** Ryan M. Layer, Fritz J. Sedlazeck, Brent S. Pedersen, Aaron R. Quinlan

## Abstract

Structural variants (SVs) are associated with cancer progression and Mendelian disorders, but challenges with estimating SV frequency remain a barrier to somatic and de novo classification. In particular, variability in filtering and variant calling heuristics limit our ability to use SV catalogs from large cohorts. We present a method to index and search the raw alignments from thousands of samples that overcomes these limitations and supports robust SV analysis.

## Main

Structural variants (SVs), which include large deletions, duplications, insertions, inversions, and translocations^1^, are increasingly associated with cancer progression and Mendelian disorders^>2–>5^. Copy number variants and gene fusions have received the most attention, but recent large-scale SV studies such as the Pan-Cancer Analysis of Whole Genomes^6^ (PCAWG), the 1000 Genome Project^7^ (1KG), gnomAD SV^8^, and the Centers for Common Disease Genomics^9^ (CCDG) have expanded our understanding of the depth and diversity of somatic SVs in cancer and polymorphic SVs in humans. Despite the importance of SVs, many barriers remain to the wide-scale adoption in disease analysis^1^. In particular, limitations to short-read SV calling, reference biases, and variability in the heuristics and filtering strategies between cohorts lead to an incomplete understanding of SV population frequency that limits our ability to assess a variant’s severity and impact^10^.

In cancer studies, SVs interpretation requires classifying variants as either germline or somatic. The gold standard is to call variants in the tumor and control tissue from the same individual together. SVs found only in the tumor are deemed somatic. This method is susceptible to the completeness and the purity of the normal sample calls, which are often sequenced at lower coverage. When a germline SV is missed in the normal tissue (e.g., due to a stochastic reduction in DNA sequence coverage), it can be incorrectly classified as somatic in chase where the variant is inherent yet missing owing to a lack of coverage. These somatic false (caused by false negatives in the control tissue) positives are wide-spread. An alternative strategy is to substitute a matched normal tissue with a panel of unrelated normal samples (e.g., 1KG, Simons Diversity Panel^11^ (SGDP)), but the time and computational costs associated with joint-calling large numbers of samples can be prohibitively high.

SV catalogs from large DNA sequencing projects (e.g., 1KG, gnomAD SV, CCDG) are used to filter tumor-only calls as a shortcut to joint calling. Variants found in both the tumor and the reference catalog can be classified as inherited since we can reasonably assume that somatic variants in general, and driver mutations in particular, are likely to be rare and unlikely to share SV breakpoints with polymorphic SVs segregating in humans. The analysis is more complicated for variants found only in the tumor calls. In principle, SVs that are not in the cohort are rare and could potentially be somatic. In practice, several SV-specific factors, including limitations in short-read calling^12^, complexities in genotyping (see **Supplementary Note 1**), and aggressive filtering to minimize false positive calls, exclude many real SVs from appearing in these reference catalogs. For example, among the thousands of cancer-related SVs that are recoverable in 1KG, an order of magnitude fewer are present in either the 1KG SV call set^7^. Given these issues, it is impossible to determine whether an SV observed in a patient and not in a reference cohort is absent from the population (i.e., a true negative) or removed in the filtering step (i.e., a false negative).

Similarly, in Mendelian disease analyses, causal variants should be either absent or are at very low-frequency in the reference population^13^. Using allele frequencies from gnomAD^14^, a catalog of single nucleotide variants (SNVs) from 141,546 human genomes, can reduce the number of variants under diagnostic consideration by two orders of magnitude^13^. Unfortunately, no equivalent resource exists for SVs since, as with the cancer analysis, static call sets from large populations are inadequate.

Pangenomes can help by identifying and genotype SVs in large populations^15^. By extending the linear reference to a graph that includes alternative haplotypes, variants, including SVs, are detected and genotyped by aligning reads directly to the sequence created by the variant. While this approach is promising, there are three major limitations. First, the standard pangenome approach to augmenting the reference involves directly adding SV sequences, which requires SV detection. As discussed above, relying on detection can be problematic. Second, graph-based methods require precise breakpoints, and in most cases short read SV-callers do not provide single-base resolution breakpoints^16^. Third, adding SVs to a pangenome is complex and including rare variants decreases performance significantly^17^. Considering these issues, pangenomes are better suited for common variants and are less useful for somatic and pathogenic variant classification.

Our solution to ensure comprehensive and accurate SV detection and allele frequency assignment is to search the raw alignments across thousands of samples using our structural variant index (STIX). For a given SV (currently deletions, duplications inversions, and translocations), STIX reports a per-sample count of every alignment that supports a given variant (**Fig. 1**). From these counts, we can, for example, conclude that a SV with high-level evidence in many healthy samples is likely a common germline variant or the product of systematic noise (e.g., reference bias), and it is unlikely to be pathogenic. By relying on the raw alignments, STIX avoids the previously described false negative issues, and recovers thousands of variants that could have otherwise been considered disease associated in seconds.

**Figure 1.**
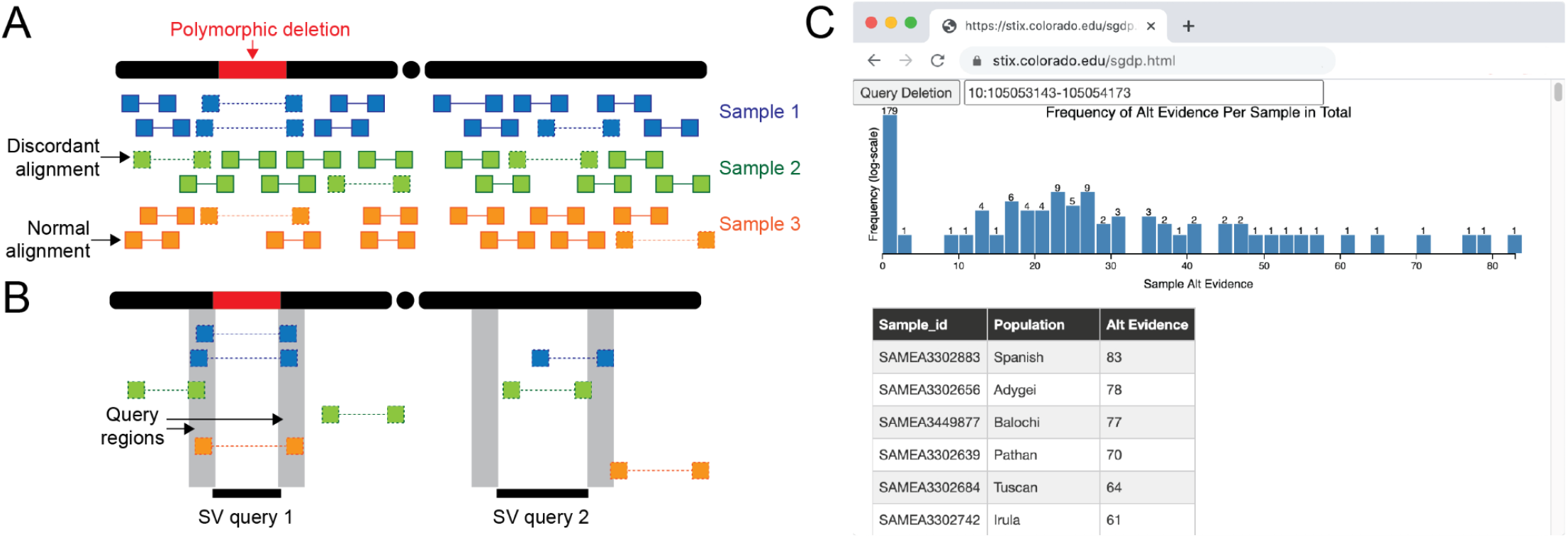
The STIX structural variant index. (**A,B**) The STIX indexing and query process for three samples and a polymorphic deletion. (**A**) Alignments tile the genome. Only a small fraction of paired end reads are discordant (designated by a dotted line connected read pairs) because of either an SV or other nonspecific causes (e.g., mapping artifacts). (**B**) Discordant alignments are extracted from all samples and indexed using GIGGLE. Query SVs are mapped to regions and alignments that reside in both regions are returned and aggregated. The first query returns three alignments in two samples and the second returns zero alignments. (**C**) The distribution of evidence depths for a deletion searched in the SGDP cohort through the http://stix.colorado.eduinterface.

STIX is built on top of the GIGGLE genome search engine^18^, which enables fast queries of thousands of genomes. Sequence alignment files (BAMs and CRAMs) contain mostly normal alignments and some (typically less than 5%) “discordant” alignments (split-reads and paired-end reads with further or shorter than expected aligned distance between pairs or an unexpected strand configuration) due to either the presence of a SV or some noise in the sequencing or alignment process (**Fig. 1A**). These alignment signals are used for detection by all current methods. STIX extracts and tracks all discordant alignments from each sample’s genome (**Figure 1A**), then creates a unified GIGGLE index for all samples. With the GIGGLE index, a user simply provides the SV type and the coordinates of its breakpoints and confidence intervals (e.g., deletion from chr10:105053143 to chr10:105054173, +/− 200), and STIX returns a sample-specific count of all alignments that support the variant. In **Figure 1B**, the first query would return two (blue reads), zero (green reads), and one (orange reads) evidence across the 3 samples. Query two would return zero evidence as no alignment supports this SV. We further have deployed web interfaces for STIX queries of 1KG and SGDP aligned to GRCh37 at http://stix.colorado.edu (**Figure 1C**). The interface takes SV coordinates as input (e.g. DEL 10:105053143-105054173) and returns the population SV evidence depth distribution in the form of a histogram and a table with the per sample SV evidence depths. The backend server that powers these interfaces also supports direct access for integrating STIX into other programs such as a VCF annotation tool.

STIX is accurate, space efficient, and fast. Considering all of the deletions, duplications, and inversions in the 1KG SV catalog, STIX identified the non-reference samples with an accuracy of 0.998, 0.995, and 0.988 respectively(see **Methods**, **Supplementary Table 1**). This result is consistent with previous reports showing STIX outperformed DELLY, SVTyper and SV2^19^. Since STIX focuses on discordant alignments, its index is a fraction of the size of the original alignments. For example, the alignments of the 1KG cohort (BAM files) required 60 terabytes of storage. The total STIX index was more than 500X smaller (110 gigabytes). Given a STIX index, population-scale queries are fast enough to scale to thousands of SVs. For example, a single SV search of the 1KG cohort using STIX completed in about 1.5 seconds while a direct extraction of the alignments alone took over 16 minutes (see **Methods**).

**Figure 2.**
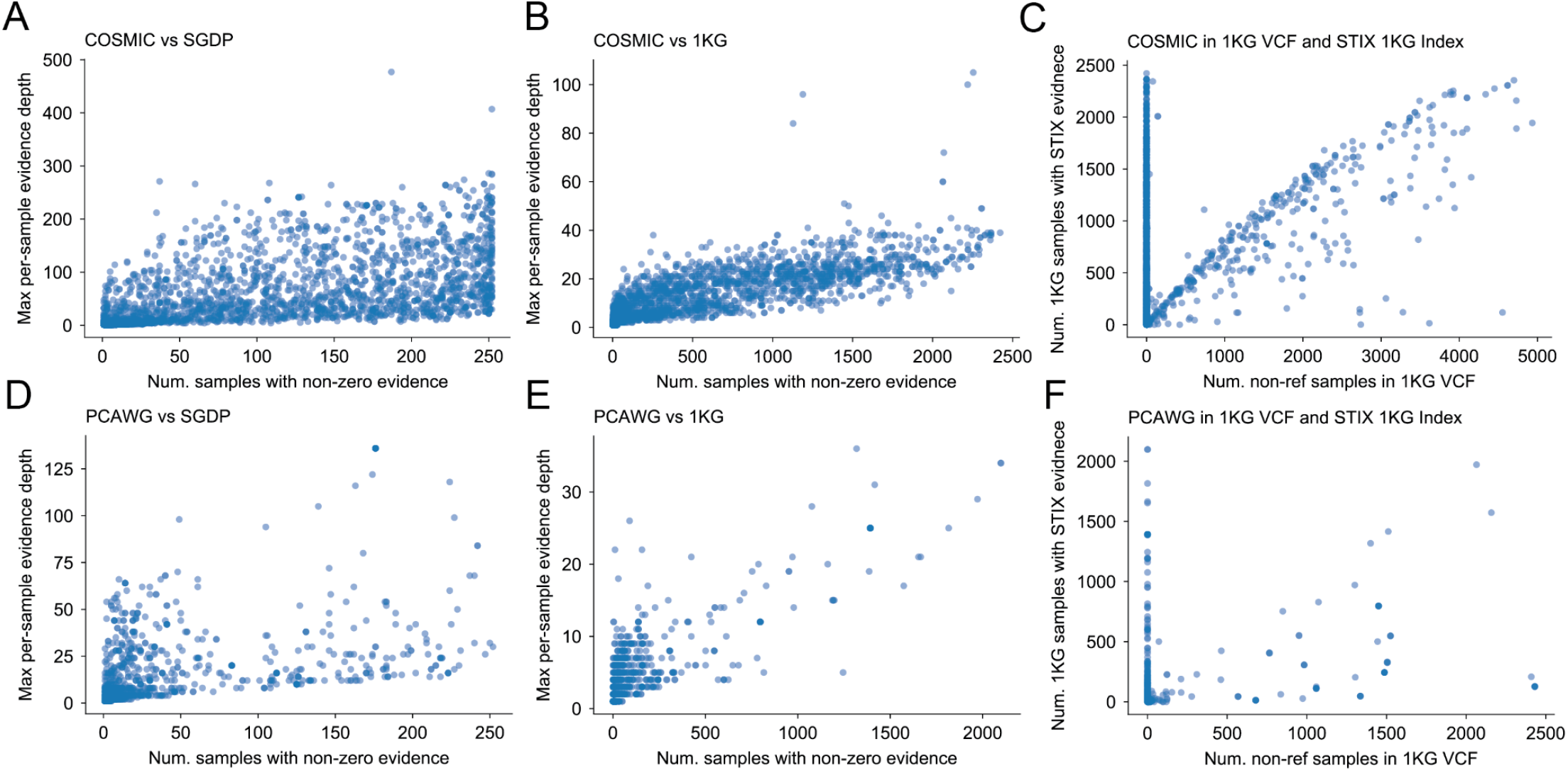
Ostensibly somatic variants appear in STIX datases. COSMIC contains 46,185 somatic deletions. Many of these SVs appear in samples from (**A**) SGDP (12,902) and (**B**) 1KG (12,2709) at high frequency and at high per-sample depth. (**C**) Only 454 COSMC SVs appear in the 1KG SV callset. (**D-F**) Similarly, PCAWG found 84,083 deletions, 2,883 of which were in SGDP and 1,732 were in 1KG. The 1KG call set contained only 193 PCAWG SVs.

Given its scalability, we can use STIX to improve somatic SV calls by scanning thousands of genomes for corroborating evidence. Among the 46,185 deletions in the Catalogue of Somatic Mutations in Cancer^20^ (COSMIC), 12,270 (26.5%) appeared in 1KG (**Fig. 2A**), 12,902 (27.9%) were in SGDP (**Fig. 2B**), and 13,295 (28.8%) were in the combined cohort (see **Supplementary Table 2**). Despite having matched normal tissues for every sample, 1,732 (2.1%) of the 84,083 somatic deletions found by PCAWG were in 1KG(**Fig. 2D**), 2,833 (3.4%) were in SGDP (**Fig. 2E**), and 3,237 (3.8%) appeared in either populations (see **Supplementary Table 3**). The SVs found by STIX are either germline or recurrent mutations and are unlikely to be driving tumor evolution. These results highlight the importance of using STIX for future studies to incorporate larger reference populations to prioritize SVs.

Scanning a large population for recurring SVs can improve somatic calling, but relying on an SV call set of the population is insufficient. While STIX found that the 12,270 COSMIC SVs had some evidence in the 1KG cohort, the published 1KG SV call set^7^ only recovered 454 variants (**Fig. 2C**). Similarly, only 193 PCAWG variants were in the 1KG catalog versus the 1,668 found by STIX (**Fig. 2F**). In both cases, many of the SVs that are missing in the catalogs were at high frequency in the populations (x=0 for **Figs. 2C** and **2F**). While the 1KG cohort is small in comparison to more recent calls sets, larger and more deeply sequenced cohorts still lag in SV recovery. For example, the gnomAD SV call set^8^ considered 27X more genomes than 1KG but only found about 2X more SVs in COSMIC and PCAWG. To fully leverage the data from these projects, we must search for evidence among the raw alignments.

In addition to somatic SVs, we used STIX to study *de novo* variation in a large family study ^21^. Since *de novo* SVs are new events, they should be rare in the population if a random process drives them. Our analysis found strong evidence (at least 3 supporting reads) for 57 of 698 de novo SVs in either 1KG or SGDP (8.7% deletions, 5.6% duplications, 30% inversions) (see **Supplementary Table 4**). Most (47) de novo SVs were observed in a single 1KG sample, and one dnSV was observed in six 1KG samples. Given the massive number of possible SV combinations (size, type, location) and the low de novo SV rate (0.16 events per genome^21^), finding any evidence in these populations highlights the plausibility of re-emerging alleles, which has been shown in other species^22^, and hotspots. Only five of the reported de novo deletions appear in the 1KG catalog. STIX again shows its utility and importance in uncovering novel insights into SV dynamics by enabling an accessible and comprehensive assessment from population data for variants often not reported in SV catalogs.

STIX enables fast and accurate SV frequency estimates directly from population-scale sequencing data, which wasn’t possible in previous SV studies due to inconsistent filtering and calling strategies. STIX overcomes these limitations by indexing all SV evidence (abnormally spaced or orientated reads) directly from the raw alignment files, avoiding detection bias, and compressing large consortia data sets down to a few gigabytes. With STIX, we indexed 2,504 samples from the 1,000 Genomes Project and 279 samples from the Simons Genome Diversity Project. These indexes helped improve somatic SV calls, highlighted the potential for recurrent de novo SVs, and are freely available for any other SV analysis at http://stix.colorado.edu. The code is freely available and open source and available at https://github.com/ryanlayer/stix.

STIX has some limitations that will be addressed in our future work, including support for insertions and long-read sequencing data and a more concise and statistically rooted population-frequency estimate. We are also exploring how STIX may enable data access with lower consent and privacy issues. By only reporting summary statistics such as population-level SV support or non-zero sample counts, the likelihood of re-identifying samples or co-occurring variants is low, which would help reconcile different consent rights across patients cohorts. With better privacy protections in place, we could pursue indexing disease genome cohorts (e.g., tumors) to help identify recurring mutations that could assist in diagnosis and treatment decisions. In any of these cases, STIX is a fast and reliable method to enable accurate genotyping of SVs across huge data collections.

## Methods

### SV evidence extraction and STIX index creation

SV alignment evidence (discordant reads and split reads) are extracted from BAM and CRAM files using excord (https://github.com/brentp/excord). Excord scans each alignment to determine if it contains a split read, has a strand configuration that is not +/−, the two aligned ends are not on the same chromosome, and the distance between the two aligned ends is further away than expected (set by the --`discordantdistance` command line parameter). The expected distance between two reads depends on the size and variance of fragments and can be measured by finding the mean and standard deviation of normal alignments in the BAM file. We recommend using the mean plus two times the standard deviation for the discordant distance. If any of these conditions is true, then the alignment is recorded as a possible piece of SV evidence. For each piece of evidence, excord stores the position and orientation of the two ends into a sample-specific BED file. For example:

|  |  |  |  |  |  |  |  |  |
| --- | --- | --- | --- | --- | --- | --- | --- | --- |
| 1 | 10022 | 10122 | 1 | 1 | 249240455 | 249240538 | 1 | 0 |
| 1 | 10031 | 10131 | 1 | 4 | 191044177 | 191044238 | 1 | 0 |
| 1 | 10036 | 10136 | 1 | 2 | 243153001 | 243153102 | -1 | 0 |
| 1 | 10054 | 10154 | -1 | 19 | 59097998 | 59098033 | -1 | 0 |
| 1 | 10066 | 10166 | -1 | 1 | 249239980 | 249240049 | -1 | 0 |

Excord was written in the Go programming language. Pre-compiled binaries are available under releases in its GitHub repository.

Each sample BED file is sorted and bgziped. For example:

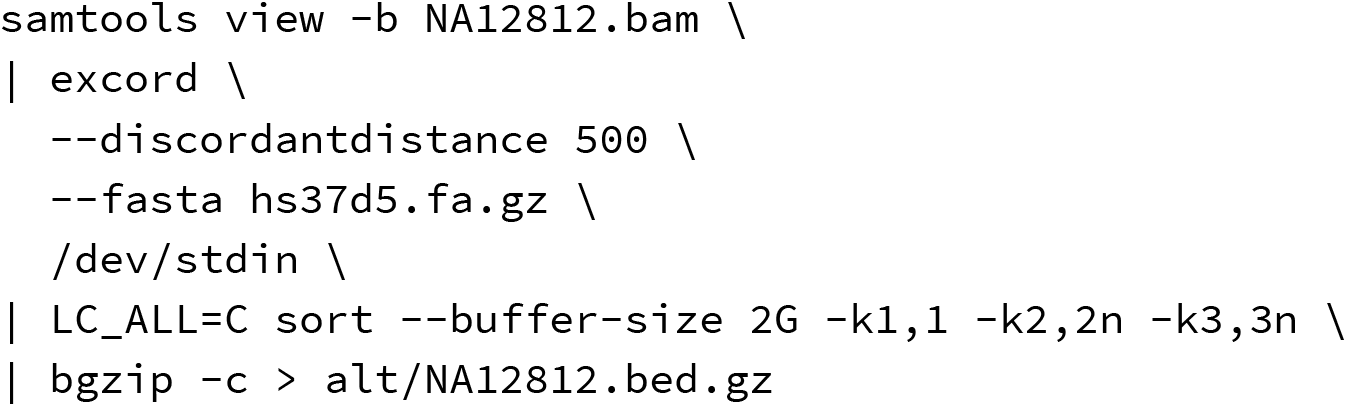

Once all sample BED files have been processed an index is created using giggle. For example:

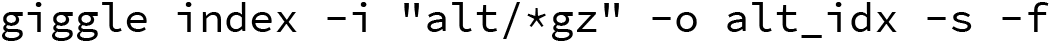

The last step is to create a sample database from a cohort pedigree file (PED). At a minimum, this file must contain a file header, and one line per sample where each line must contain the sample name and the name of its associated BED file. The following example has three extra fields:

| SampleSex | Population | Super_Population | Alt_File |  |
| --- | --- | --- | --- | --- |
| NA12812 | 1 | CEU | EUR | NA12812.bed.gz |
| HG00672 | 2 | CHS | EAS | HG00672.bed.gz |
| NA12878 | 2 | CEU | EUR | NA12878.bed.gz |
| HG00674 | 1 | CHS | EAS | HG00674.bed.gz |

Creating the sample database requires the giggle index, input PED file name, output database name, and the column number that contains the name of the sample BED file. For example:

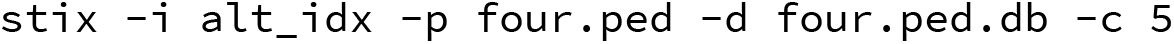

Once the BED files have been indexed and the sample database has been created from the PED file, STIX can now query the samples for SV evidence. For each query, the user must specify the index location (-`i`), sample database (-`d`), SV type (-`t`),left (-`l`) and right (-`r`) breakpoint coordinates, and window size (-`s`) to consider around each breakpoint. The window size will depend on the size and variance of the sample fragments. We recommend using the same value used for the discordant distance parameter in the excord extraction. The output of STIX is a per-sample count of alignments that support the existence of the SV in the sample. For example:

| Id | Sample | Sex | Population | Super_Population | Alt_File | Pairend | Split |
| --- | --- | --- | --- | --- | --- | --- | --- |
| 0 | HG00672 | 2 | CHS | EAS | HG00672.13.14.bed.gz | 8 | 0 |
| 1 | HG00674 | 1 | CHS | EAS | HG00674.13.14.bed.gz | 7 | 0 |
| 2 | NA12812 | 1 | CEU | EUR | NA12812.13.14.bed.gz | 7 | 0 |
| 3 | NA12878 | 2 | CEU | EUR | NA12878.13.14.bed.gz | 11 | 0 |

### 1,000 genomes phase three STIX index

2,504 low-coverage BAMs (GRCh37) and the PED file were downloaded from the 1,000 genomes AWS S3 bucket (`s3://1000genomes/phase3/data/`). Excord was run on each sample with --`discordantdistance` set to 500.

### Simons Genome Diversity Panel STIX index

252 30X-coverage FASTQ files and PED file were downloaded from the Simons Foundation (https://www.simonsfoundation.org/simons-genome-diversity-project/) and aligned to the human reference genome (GRCh37) using BWA-MEM. Excord was run on each sample with --`discordantdistance` set to 500.

### STIX speed measurement

To test the speed of STIX versus any other alternative genotyping method that accesses the BAMs directly, we compared the time required for STIX to query a specific SV (DEL, 10:105053143-105054173) across the full 1KG cohort versus how much time was required to read the alignments in the same region of each BAM in the 1KG cohort. The assumption being that any genotyping method would need to at least read the alignments, and the I/O time would be a lower bound for any such method.

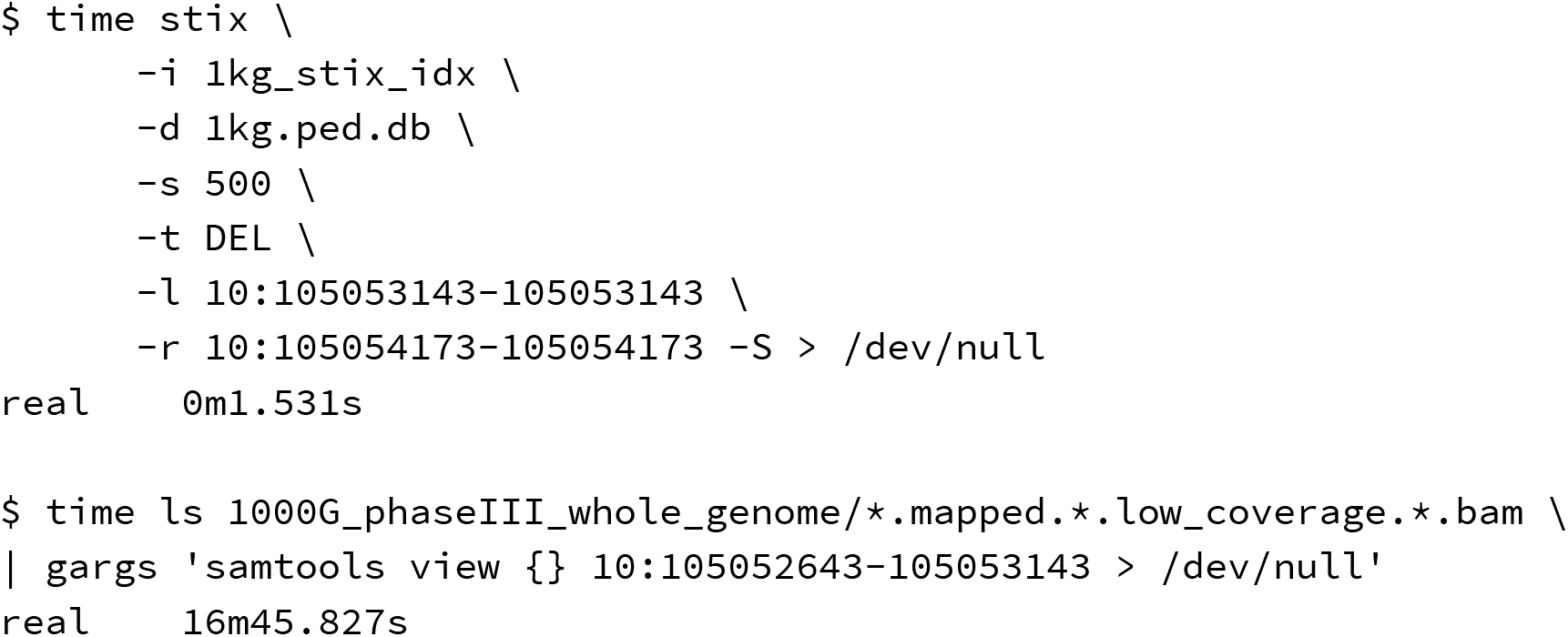

### Accuracy measurement

To determine STIX’s classification performance we considered the 1KG cohort and the phase 3 SVs identified by Sudmant et al.^7^. For each reported deletion, duplication, and inversion, we collected the samples that were identified by 1KG as being non-reference. This analysis only included SVs with the CIEND and CIPOS values specified.

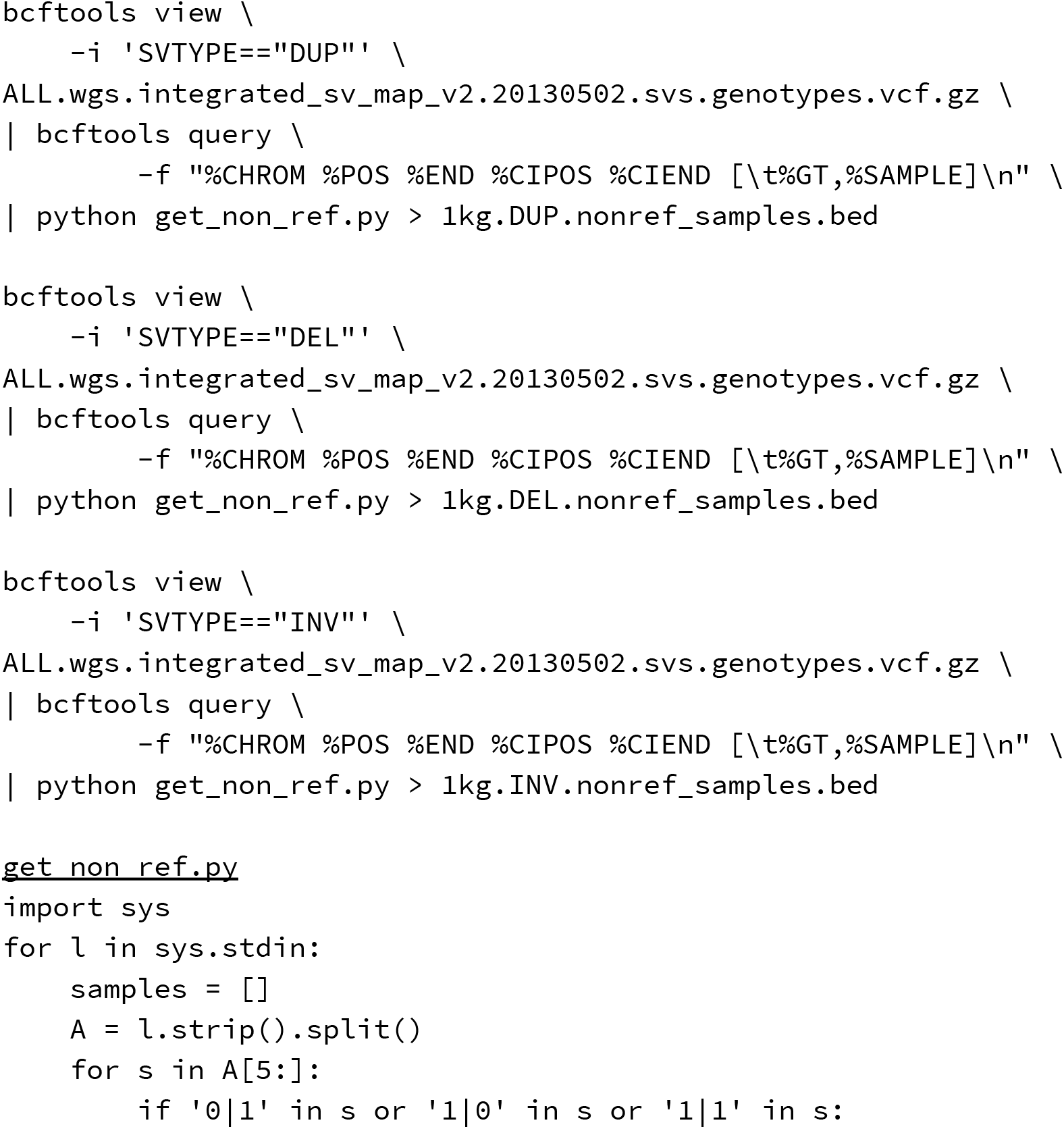
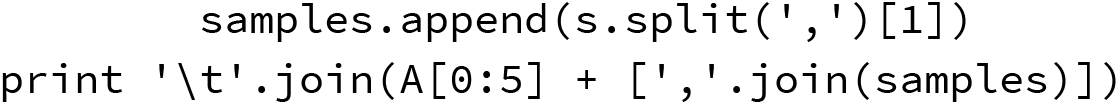

For each of those SVs, we then constructed a similar list of samples where STIX found evidence of the same variant.

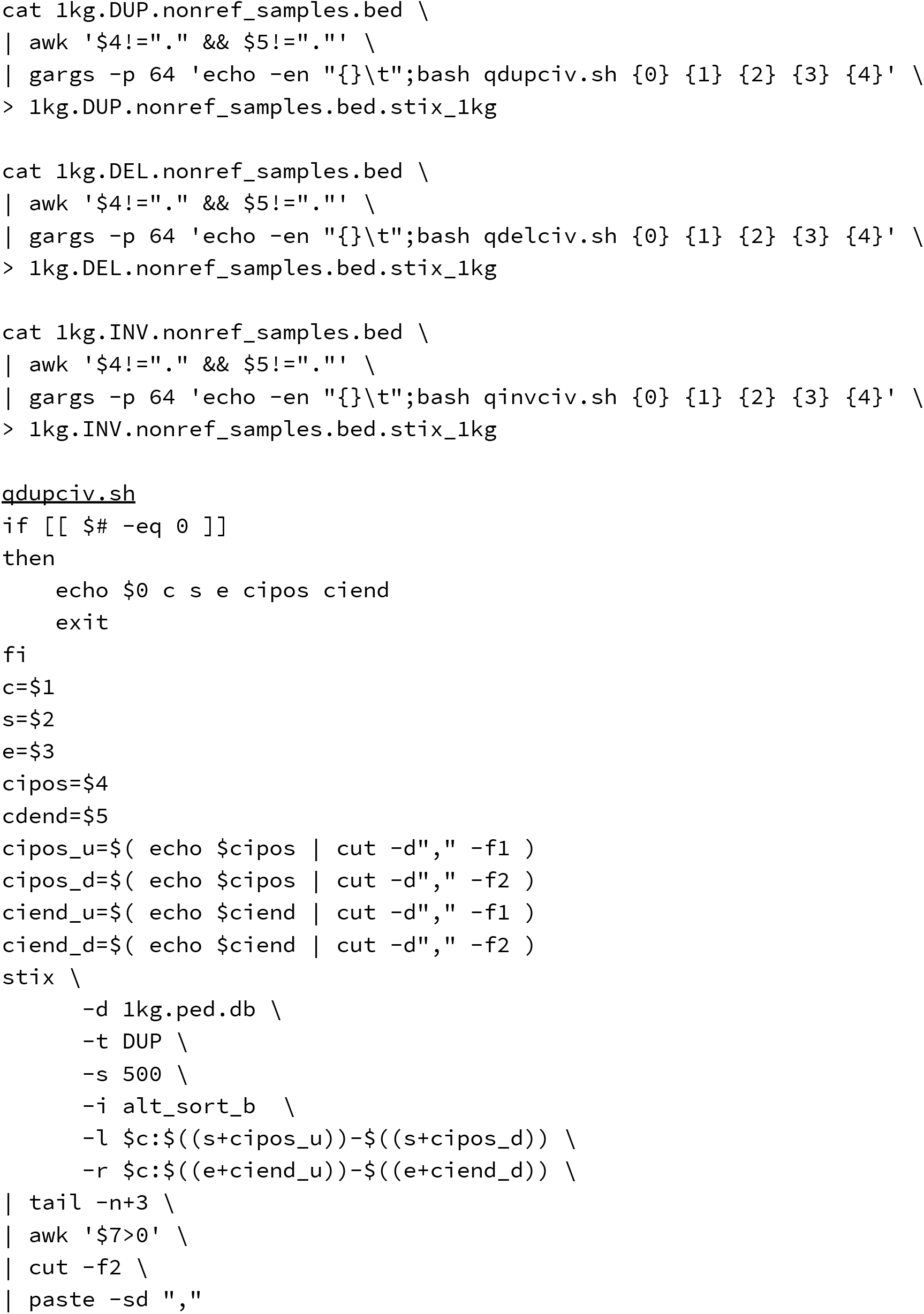

Given the list of non-reference samples from the 1KG catalog and the list of samples with supporting evidence from STIX, we computed the following values for deletions, duplications, and inversions separately.

- positives (P): Number of non-reference samples in the 1KG catalog
- negatives (N): Number of reference samples in the 1KG catalog (total samples minus positives)
- true positives (TP): Number of samples with evidence from STIX that were non-reference in the 1KG catalog
- true negatives (TN): Number of samples with no evidence from STIX that were reference in the 1KG catalog
- false positives (FP): Number of samples with evidence from STIX that were reference in the 1KG catalog
- false negatives (FN): Number of samples with no evidence from STIX that were non-reference in the 1KG catalog

From these values we computed:

- accuracy = (TP + TN) / (P+N)
- precision = TP/(TP + FP)
- sensitivity = TP/P
- specificity = TN/N
- F1 = 2TP/(2TP+FP+FN)

### COSMIC SV evaluation

The COSMIC SV catalog was downloaded from the COSMIC website (https://cancer.sanger.ac.uk/cosmic/download, Structural Genomic Rearrangements, login required). The chromosomal position of the deletions (`intrachromosomal deletion`), duplications (`intrachromosomal tandem duplication`), and inversions (`intrachromosomal inversion`) were extracted and sorted into a compressed BED file.

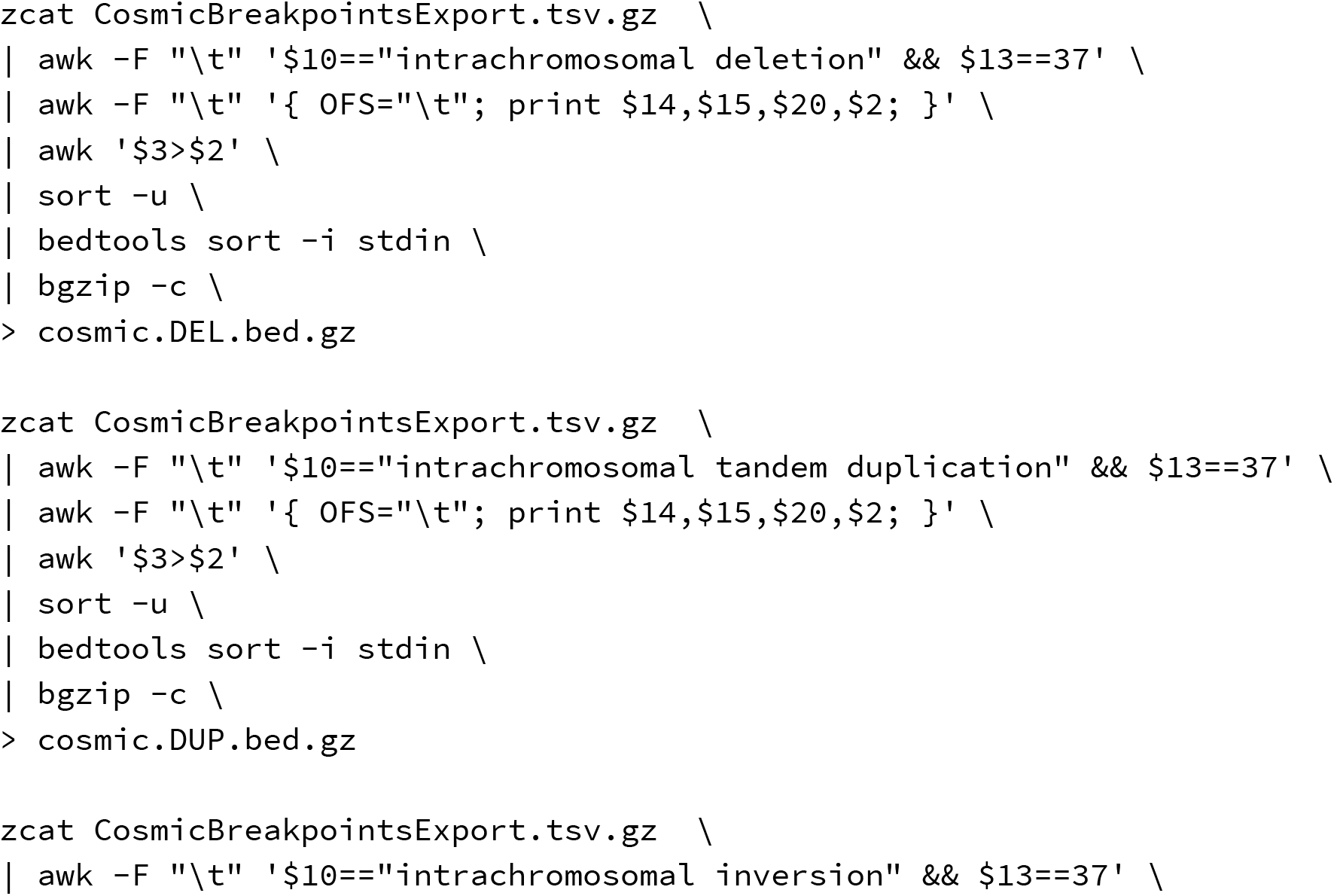
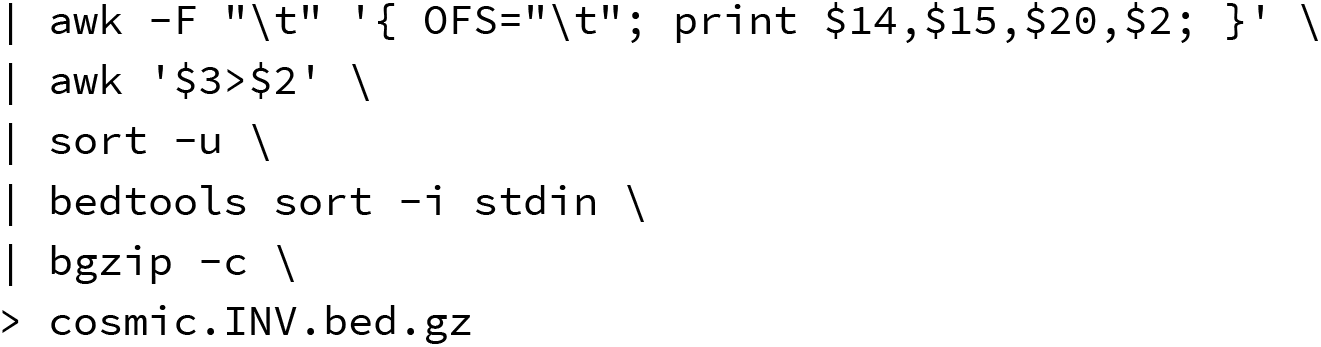

To determine the overlap between the COSMIC SVs and the 1KG catalog, we converted the 1KG SV VCF to SV-type specific BED files and intersected these files with the corresponding COSMIC BED files. Intersections required a reciprocal overlap of 90%. From these intersections we compute the 1KG allele frequency.

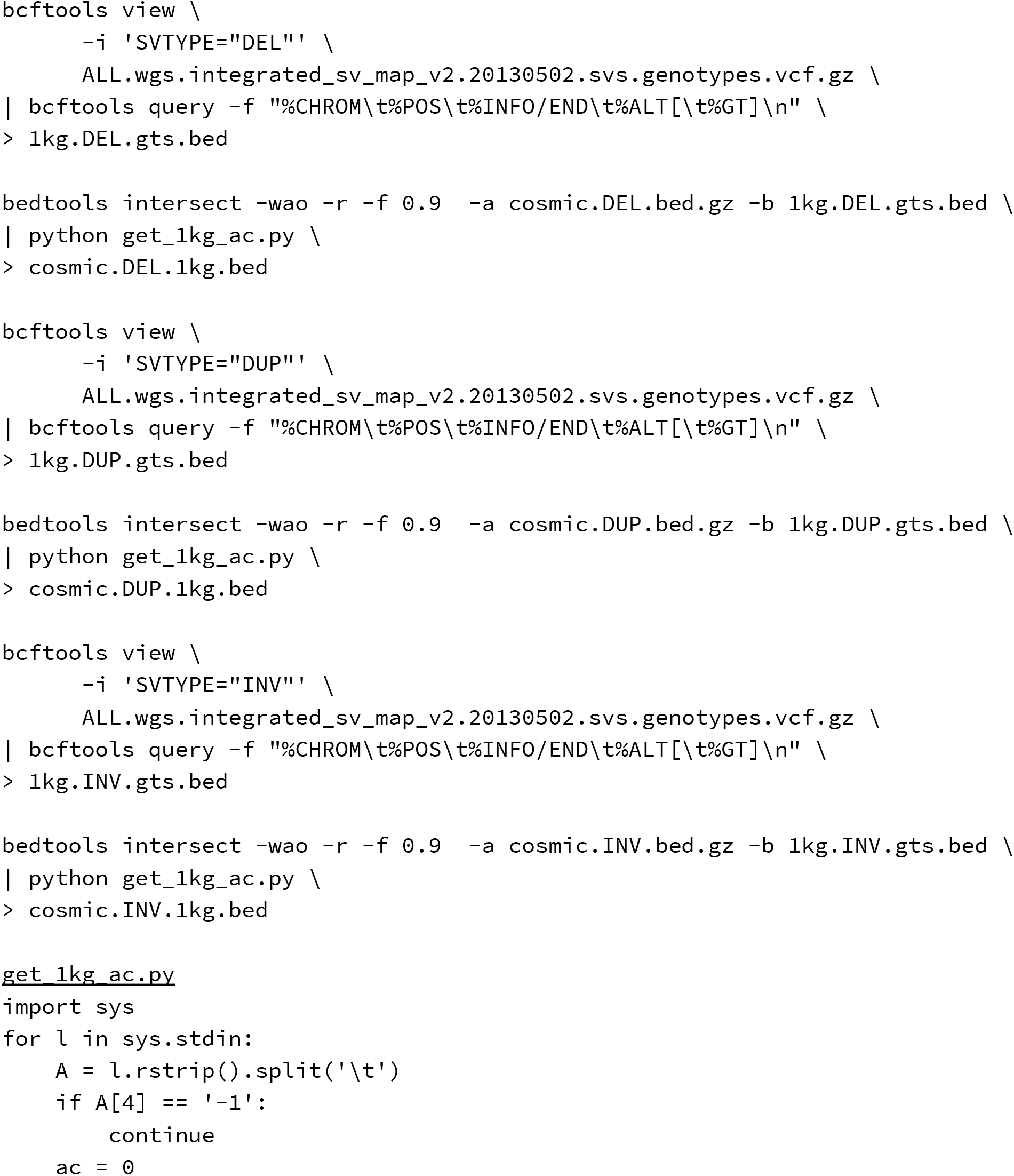
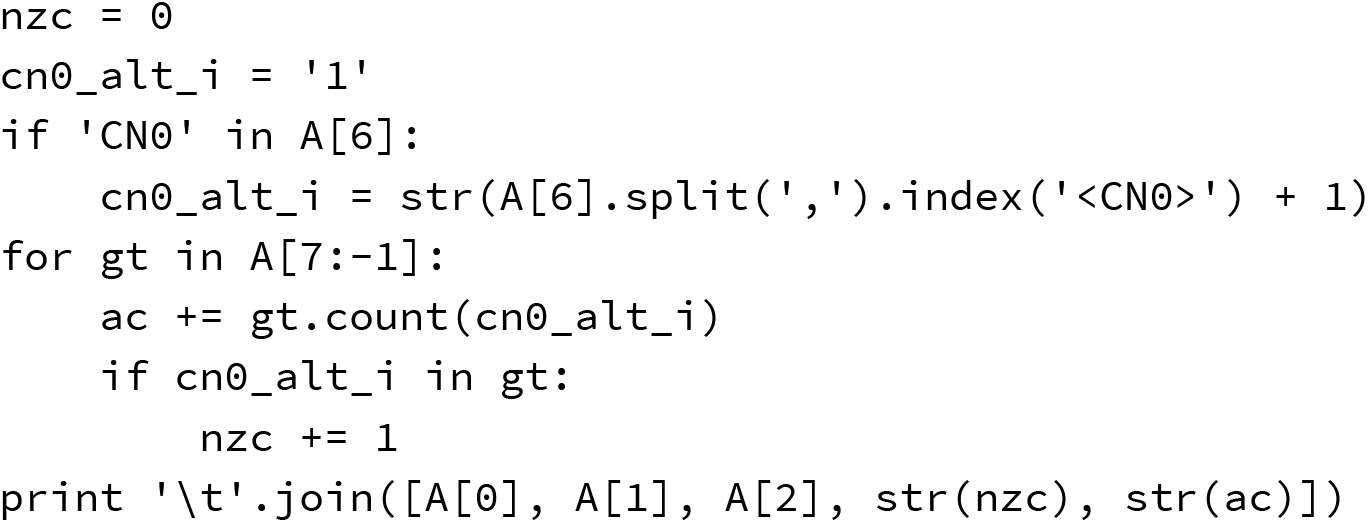

To determine the overlap between the COSMIC SVs and the gnomAD SV catalog, we retrieved the version 2.1 SV BED file from the gnomAD web site (https://gnomad.broadinstitute.org/downloads/#v2-structural-variants) and split the BED file into SV-type specific BED files and intersected these files with the corresponding COSMIC BED files. Intersections required a reciprocal overlap of 90%.

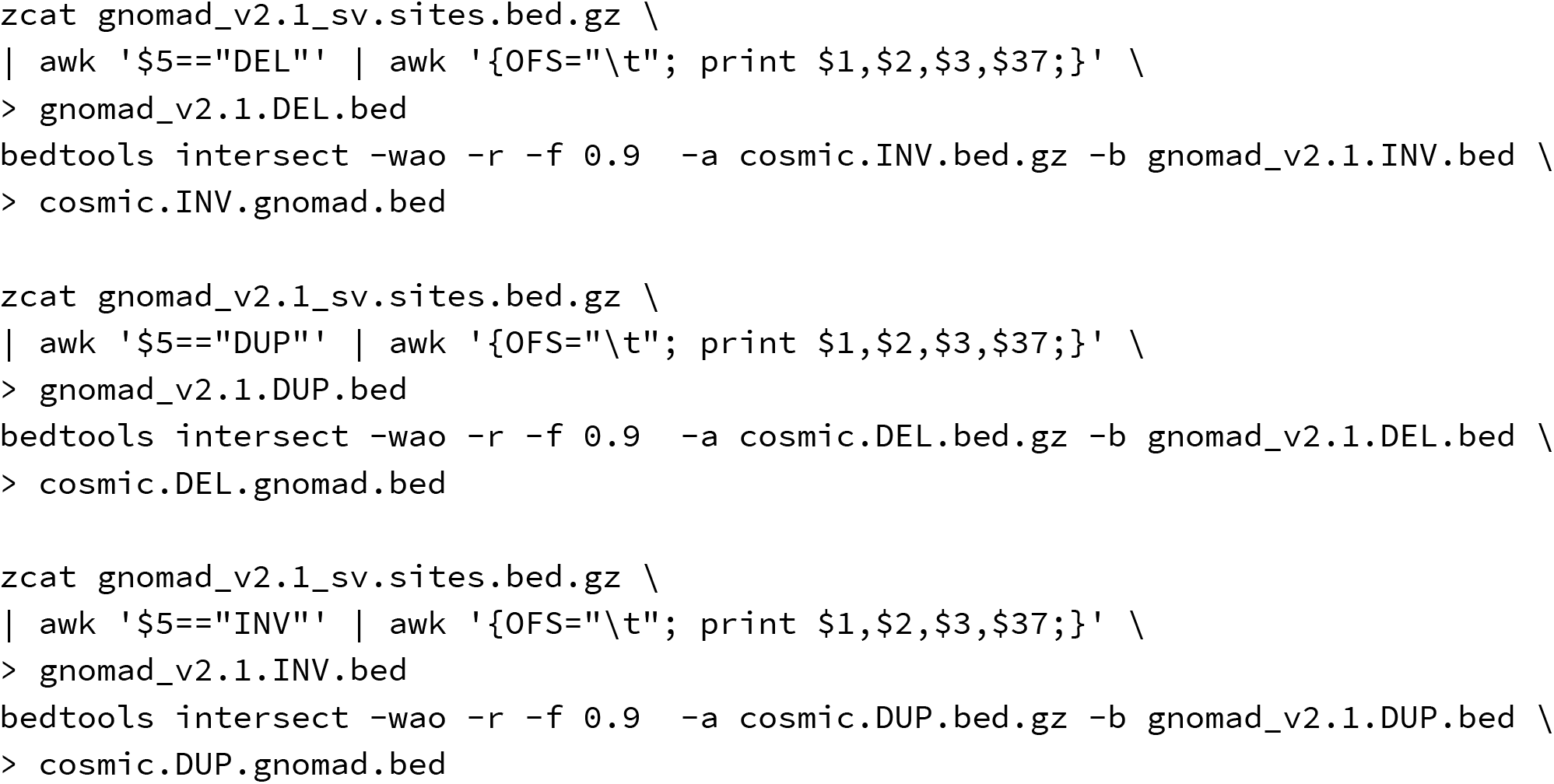

To determine the overlap between the COSMIC SVs and the STIX for 1KG and SGDP, we submitted a STIX query for each SV in the COSMIC SV-type BED files using a 500 base pair window. For each SV we compute the number of samples with some supporting evidence.

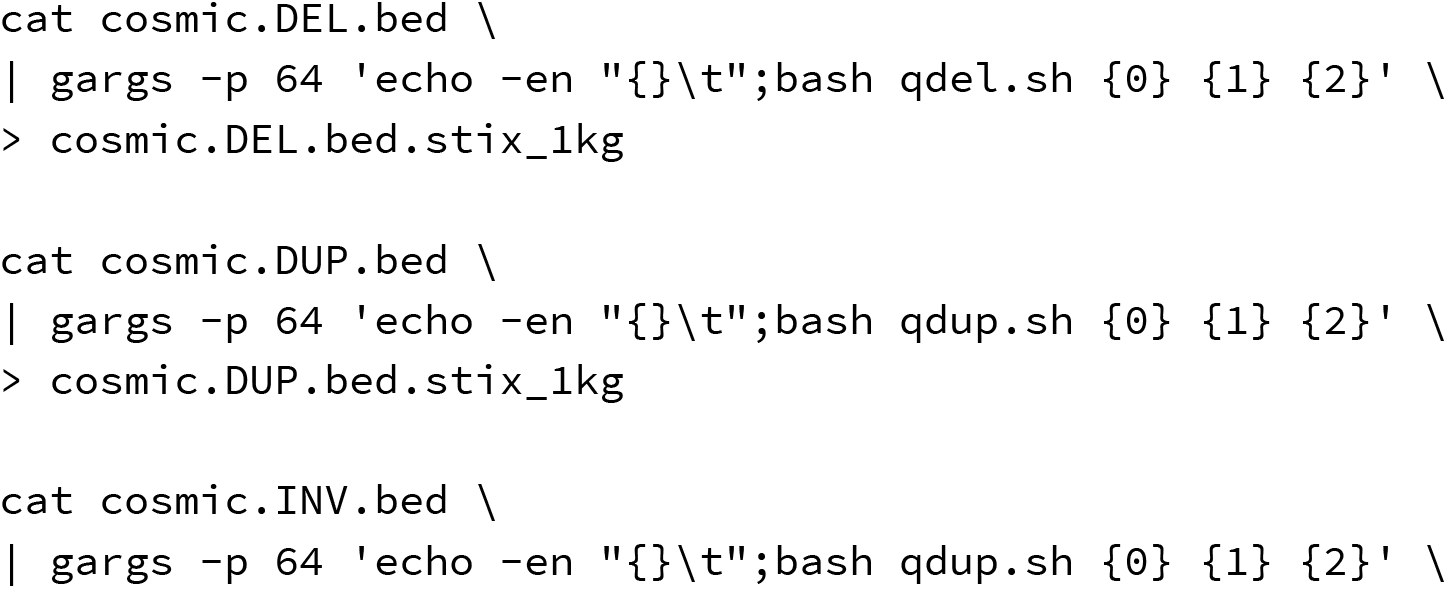
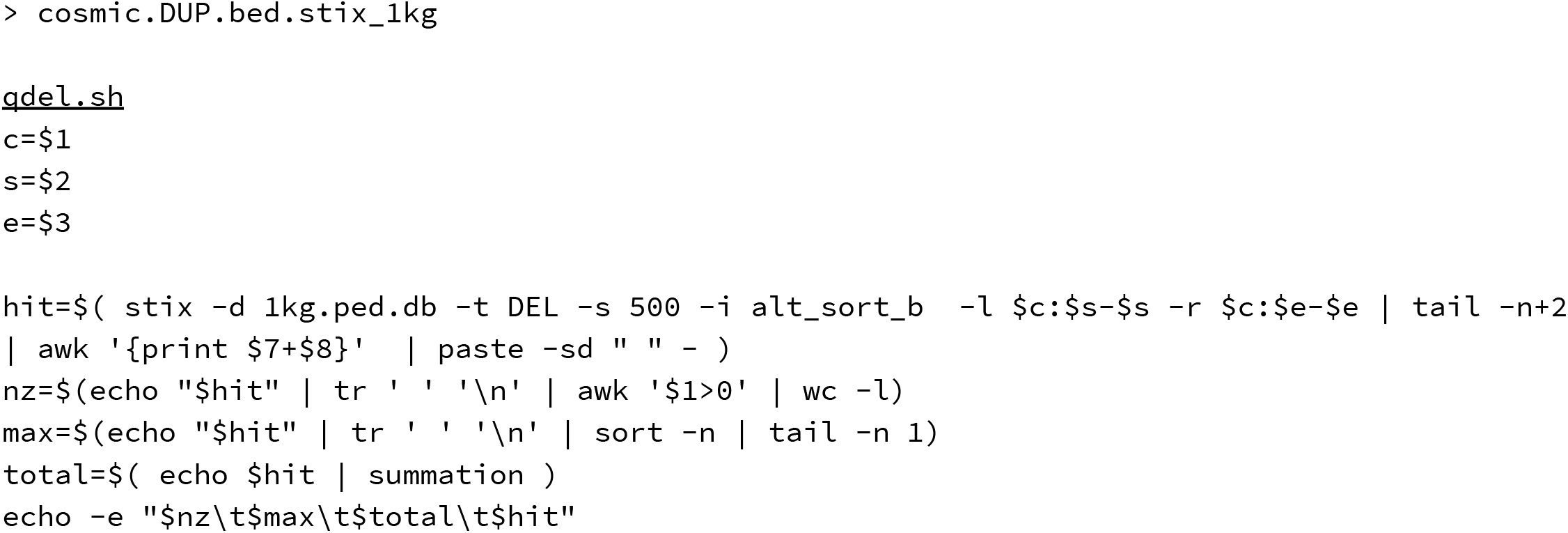

### Pan-Cancer Analysis of Whole Genomes SV evaluation

The PCAWG SV catalogs were downloaded from the ICGC data portal website (https://dcc.icgc.org/releases/PCAWG/consensus_sv/), and combined SV-type specific call sets.

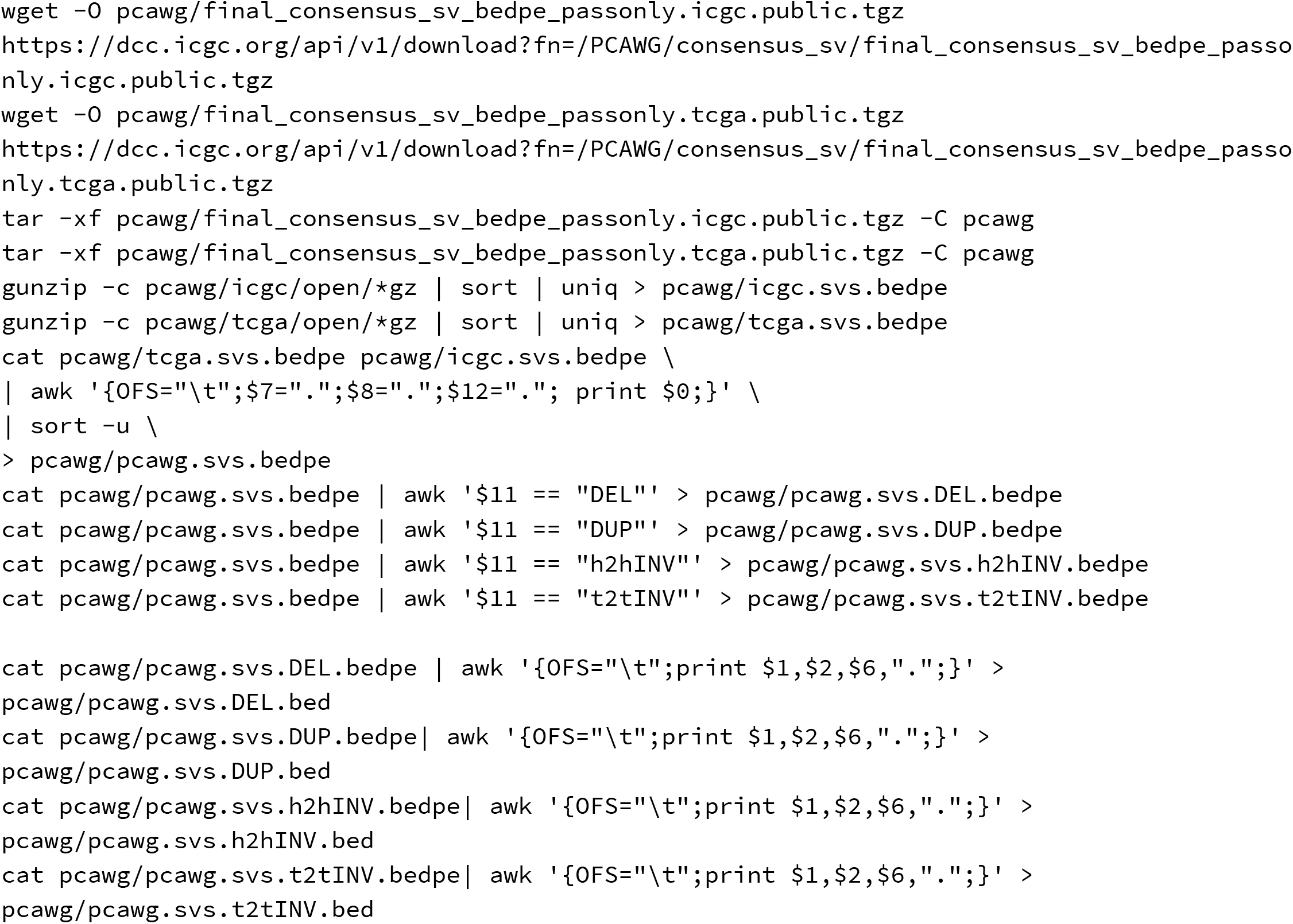

Similar to the process in *COSMIC SV evaluation, to* determine the overlap between the PCAWG SVs and the 1KG catalog, we converted the 1KG SV VCF to SV-type specific BED files and intersected these files with the corresponding PCAWG BED files. Intersections required a reciprocal overlap of 90%. From these intersections we compute the 1KG allele frequency.

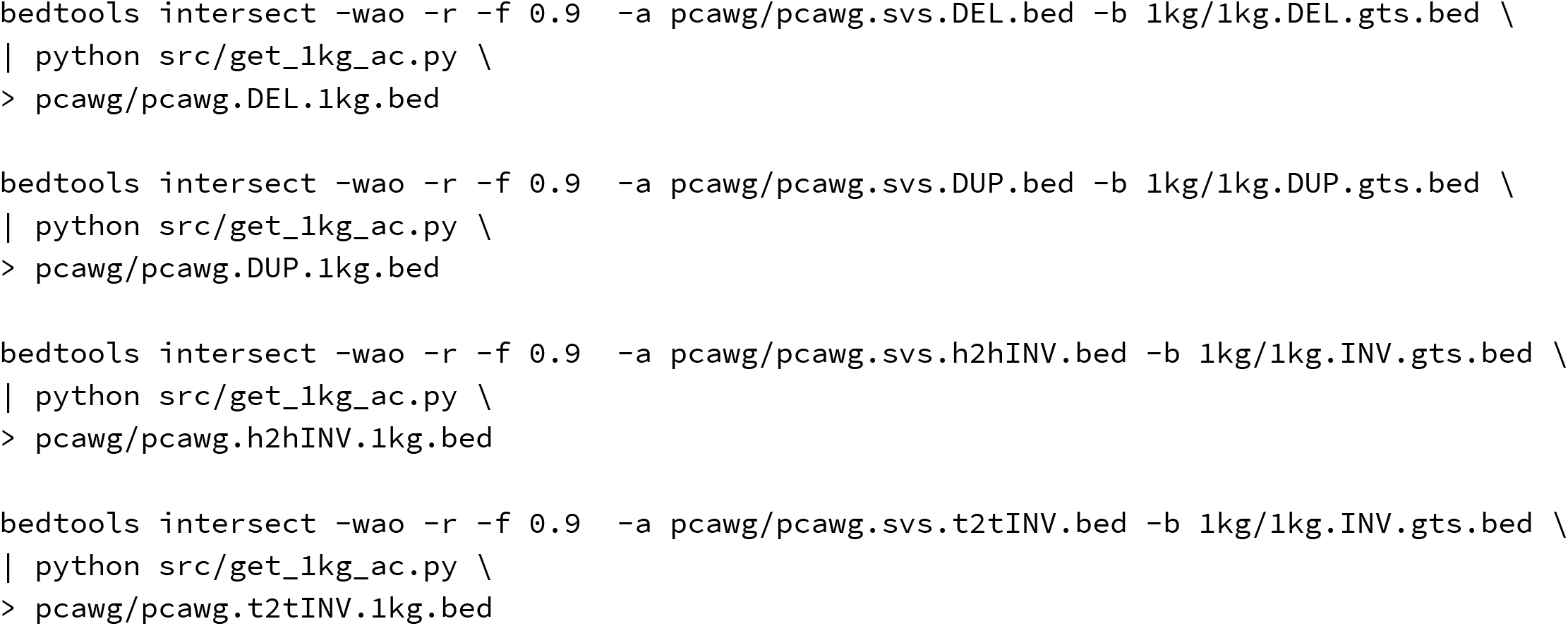

To determine the overlap between the PCAWG SVs and the gnomAD SV catalog, we retrieved the version 2.1 SV BED file from the gnomAD web site (https://gnomad.broadinstitute.org/downloads/#v2-structural-variants) and split the BED file into SV-type specific BED files and intersected these files with the corresponding PCAWG BED files. Intersections required a reciprocal overlap of 90%.

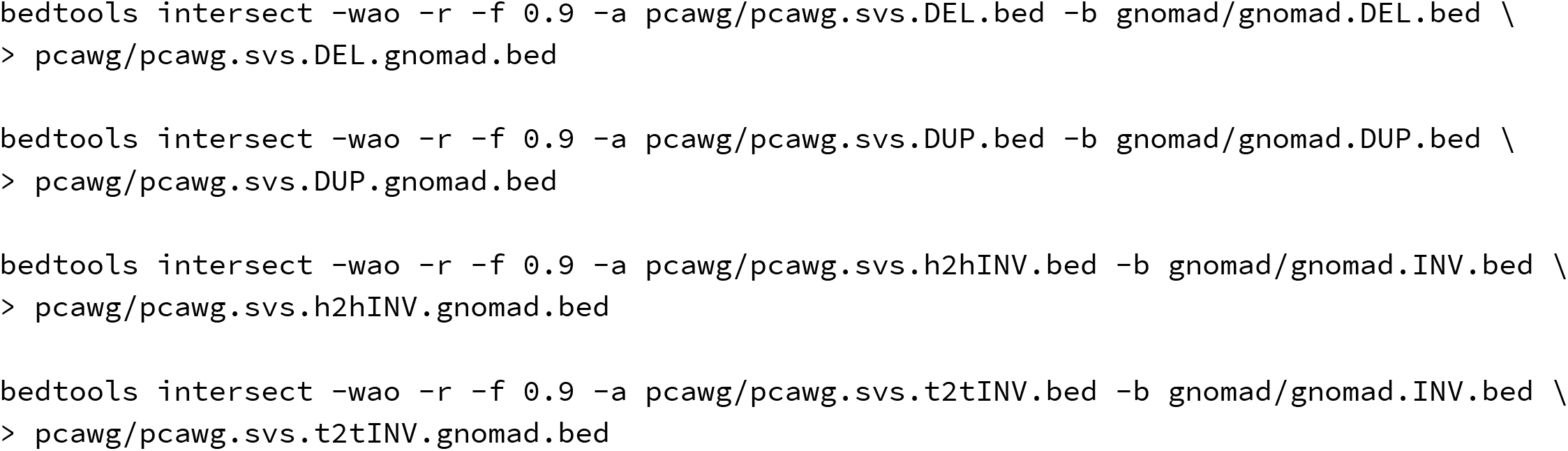

To determine the overlap between the PCAWG SVs and the STIX for 1KG and SGDP, we submitted a STIX query for each SV in the PCAWG SV-type BEDPE files using a 500 base pair window. For each SV we compute the number of samples with some supporting evidence.

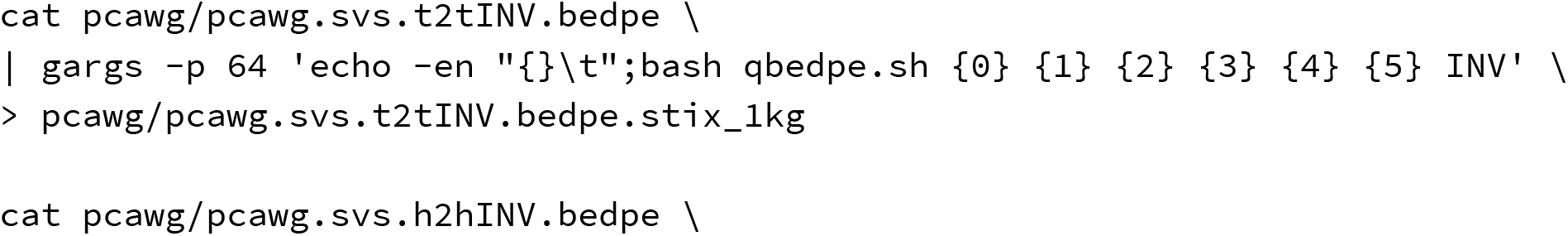
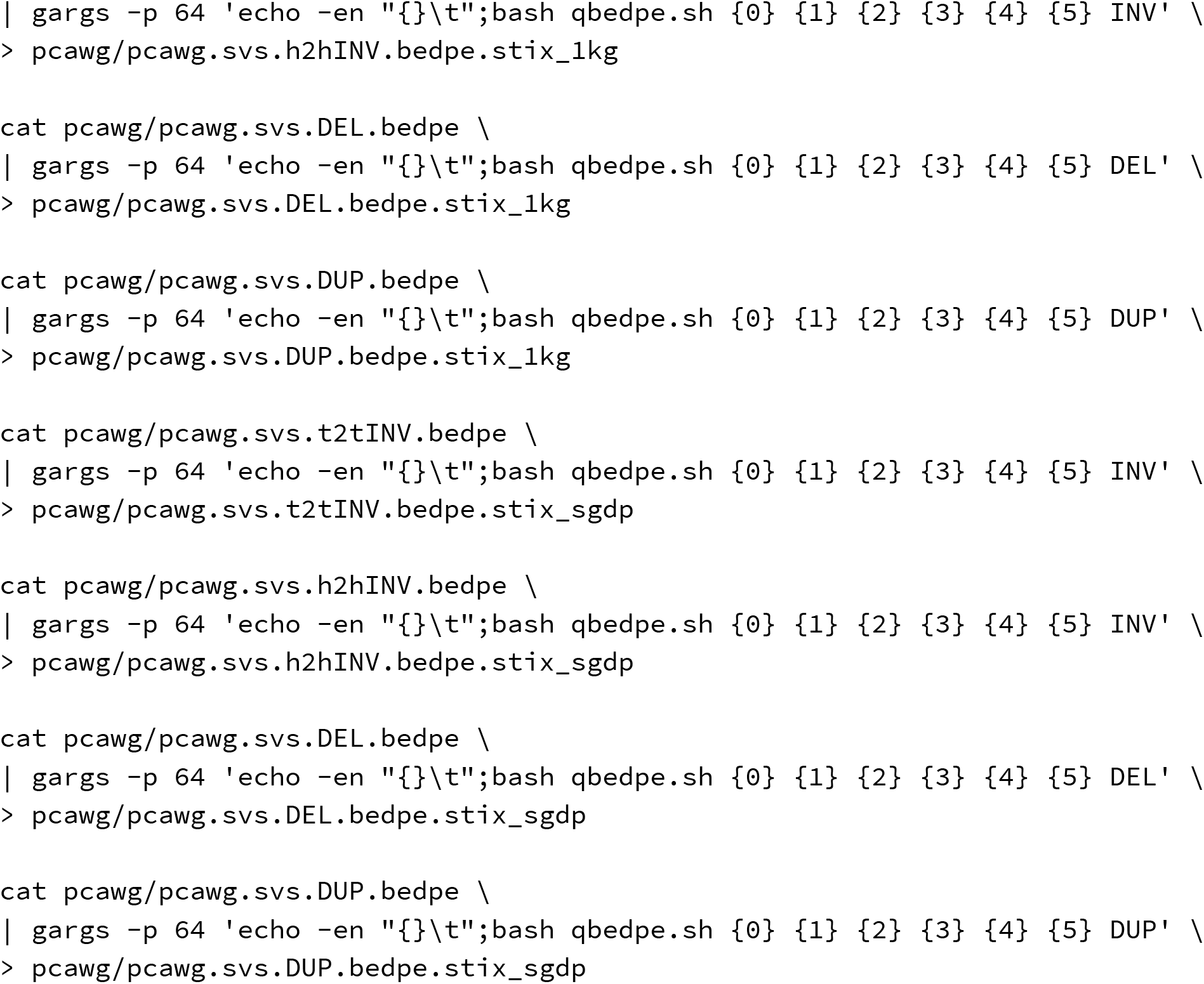

### De novo SV evaluation

The de novo SV catalog was retrieved from the GitHub repository referenced in the publication. Those SVs were reported using the GRCh38 human reference genome. We used the UCSC genome browser tools to lift the SVs to GRCH37, then split the file into SV type specific BED files.

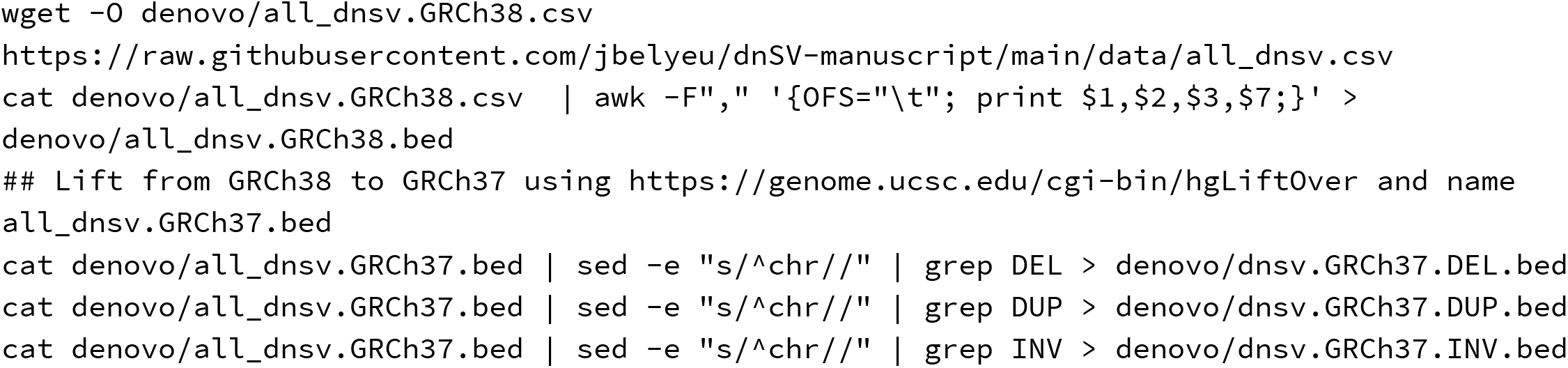

Similar to the process in *COSMIC SV evaluation, to* determine the overlap between the de novo SVs and the 1KG catalog, we converted the 1KG SV VCF to SV-type specific BED files and intersected these files with the corresponding PCAWG BED files. Intersections required a reciprocal overlap of 90%. From these intersections we compute the 1KG allele frequency.

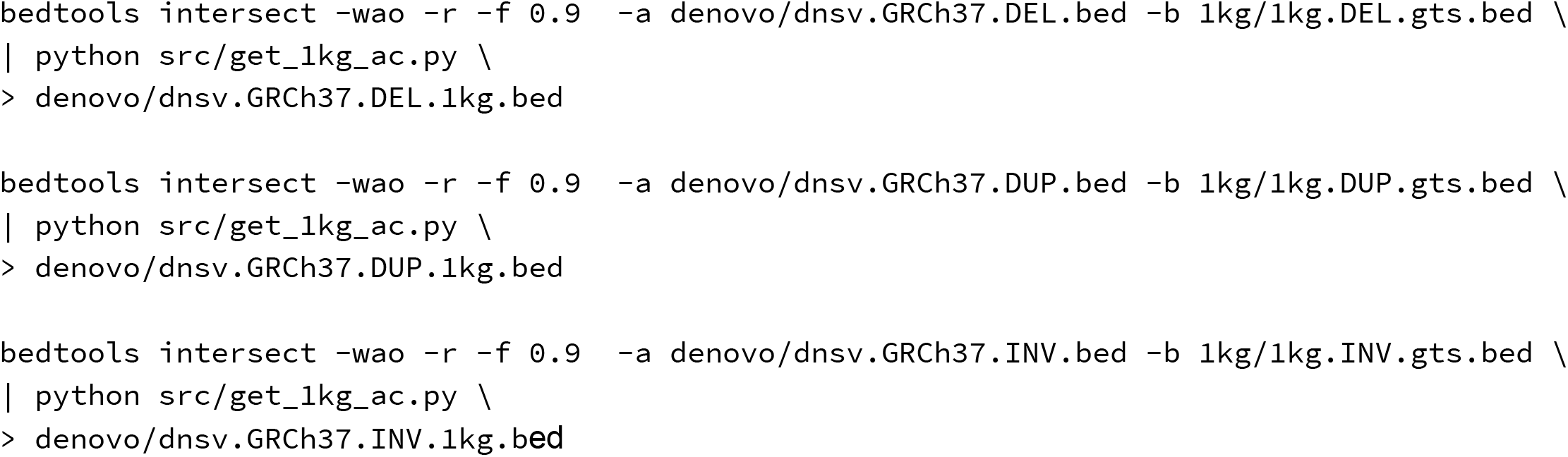

To determine the overlap between the de novo SVs and the gnomAD SV catalog, we retrieved the version 2.1 SV BED file from the gnomAD web site (https://gnomad.broadinstitute.org/downloads/#v2-structural-variants) and split the BED file into SV-type specific BED files and intersected these files with the corresponding PCAWG BED files. Intersections required a reciprocal overlap of 90%.

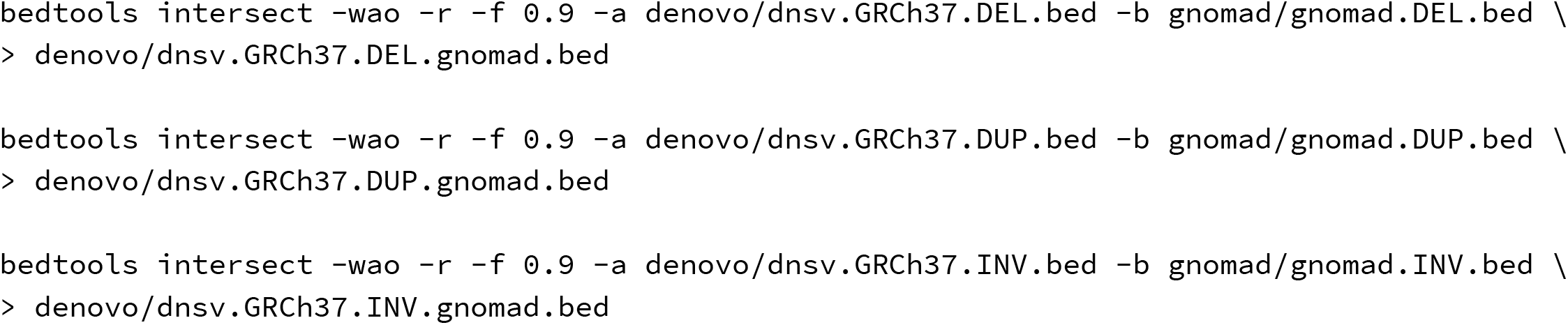

To determine the overlap between the de novo SVs and the STIX for 1KG and SGDP, we submitted a STIX query for each SV in the de novo SV-type BED files using a 500 base pair window. For each SV we compute the number of samples with some supporting evidence.

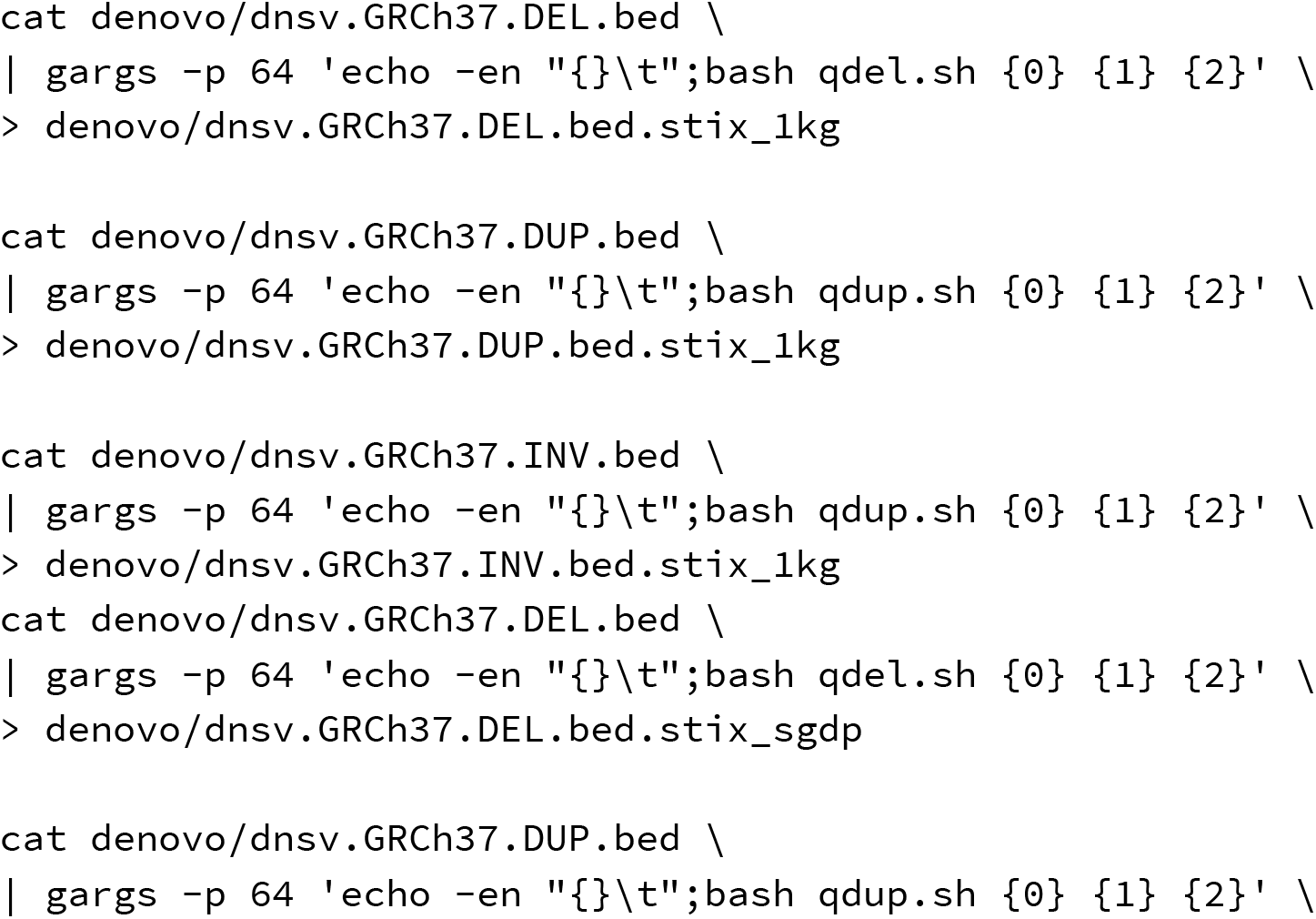
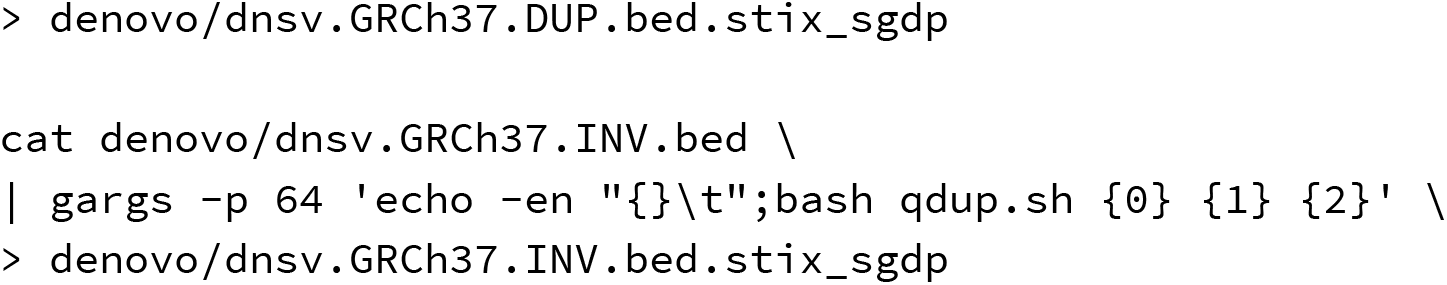

## Supplementary Notes

1. Challenges with calling and genotyping structural variants

**Supplement Note Figure S1.**
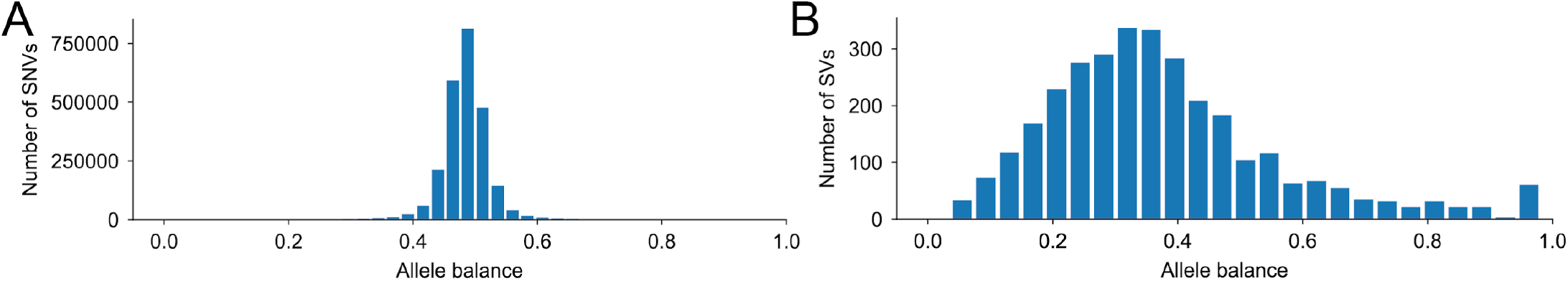
Allele balance (number of reads matching the reference/total reads) for (**A**) SNVs and (**B**) SVs for the HG002 individual from the Genome in a Bottle Consortium.

Part of the challenge when moving from SNVs to SVs is the substantial increase in the uncertainty of the underlying data. For example, the allele balance for heterozygous SNVs and SVs from the Genome In a Bottle Consortium^23,24^ sample shows a shift from the expected peak at 0.5 allele balance in SNVs (**Fig. 1A**) to 0.3 in SVs (**Fig. 1B**). The reason for this shift is that SV detection and genotyping from short-read data is complicated by evidence that does not provide direct information about the location of the variant (e.g., read depth and discordant pair-reads). These two issues result in fundamentally different detection and genotyping strategies for SVs. Instead of explicitly testing for the existence of every possible SV (which is intractable), read alignment evidence is clustered, and a consensus breakpoint (which is often not at single-base resolution) and genotype is inferred. The two major issues with this type of clustering are instances where spurious alignments overlap by chance, causing false positives, and where fluctuations in coverage create false negatives or incorrect genotypes. Both of these cases produce SVs with a wide range of per-sample evidence depths and summarizing each sample into just three states (homozygous reference, heterozygous, and homozygous alternate) hides information that can be important when determining if a newly observed variant is common, rare, or noise. Genotype quality scores capture some of this uncertainty, but in practice, these scores are only used to exclude problematic samples from an analysis. This highlights the need for new metrics that can represent the full extent of structural variant evidence in a population.

### Supplementary Tables

**Supplementary Table 1.** STIS performance across SV types considering the 1KG SV calls. In general, STIX performed well across all SV types and did exceptionally well for accuracy, precision, and specificity. The one exception was that STIX had a high number of false-positive duplication calls, leading to low precision and sensitivity. Upon inspection, just seven loci accounted for 95% of the false-positive calls. For these duplications, STIX estimated a much higher population frequency than what was listed in the 1KG catalog.

|  | Deletions | Duplications | Inversions |
| --- | --- | --- | --- |
| accuracy | 0.989 | 0.995 | 0.988 |
| precision | 0.955 | 0.135 | 0.962 |
| sensitivity | 0.645 | 0.514 | 0.713 |
| specificity | 0.999 | 0.996 | 0.999 |
| F1 | 0.770 | 0.213 | 0.819 |

**Supplementary Table 2.**
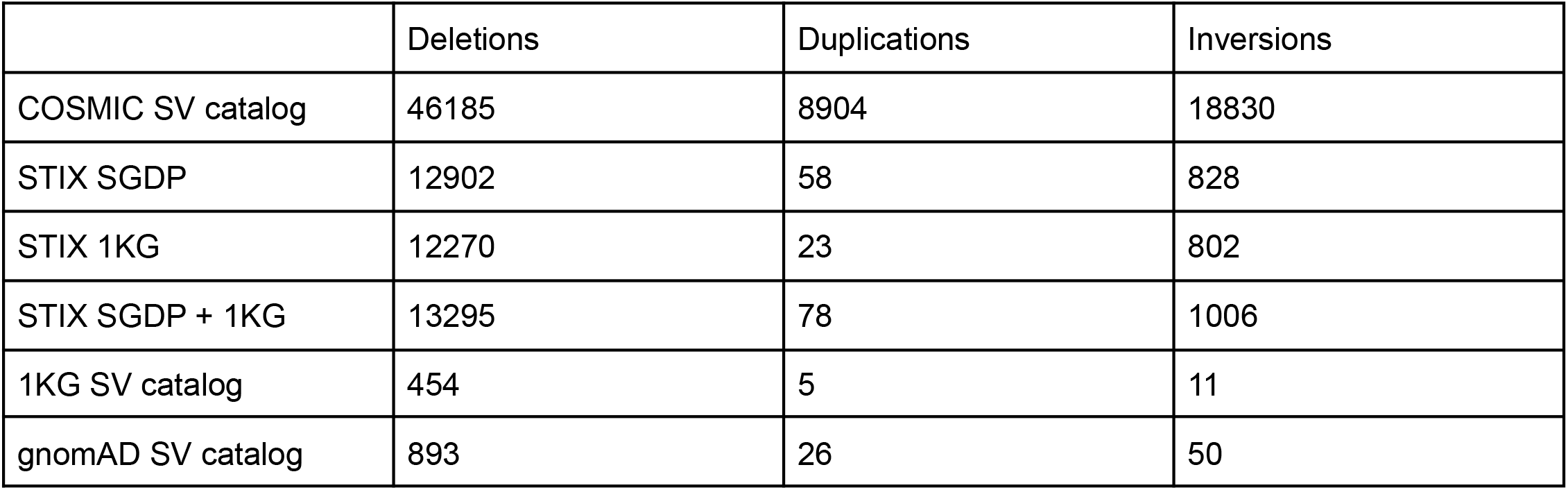
The frequency of purportedly somatic SVs from the COSMIC database considering different SV collections. STIX consistently found evidence for many more COSMIC SVs than other sources even when considering the same underlying samples (i.e., STIX 1KG versus the 1KG SV catalog).

**Supplementary Table 3.** The frequency of purportedly somatic SVs identified by the PCAWG study considering different SV collections.

|  | Deletions | Duplications | H2H Inversions | T2T Inversions |
| --- | --- | --- | --- | --- |
| PCAWG catalog | 84083 | 72764 | 38602 | 37613 |
| STIX SGDP | 2833 | 790 | 3091 | 2893 |
| STIX 1KG | 1732 | 221 | 2838 | 2641 |
| STIX SGDP + 1KG | 3237 | 843 | 3531 | 3284 |
| 1KG catalog | 193 | 40 | 1 | 1 |
| gnomAD catalog | 433 | 165 | 88 | 101 |

**Supplementary Table 4.**
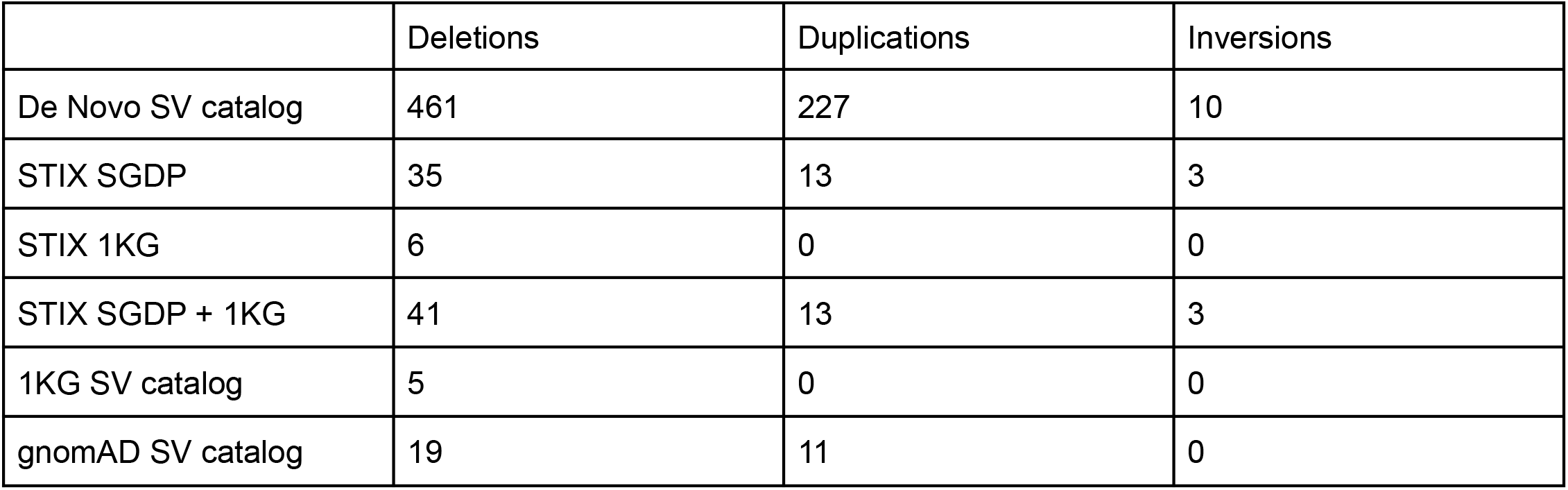
The frequency of purportedly de novo SVs from a large family study. For the STIX counts, samples had at least three supporting reads.

|  | Deletions | Duplications | Inversions |
| --- | --- | --- | --- |
| De Novo SV catalog | 461 | 227 | 10 |
| STIX SGDP | 35 | 13 | 3 |
| STIX 1KG | 6 | 0 | 0 |
| STIX SGDP + 1KG | 41 | 13 | 3 |
| 1KG SV catalog | 5 | 0 | 0 |
| gnomAD SV catalog | 19 | 11 | 0 |

## Notes

https://stix.colorado.edu/

https://github.com/ryanlayer/stix

### Competing Interest Statement

The authors have declared no competing interest.

